# Opioid Suppression of an Excitatory Pontomedullary Respiratory Circuit by Convergent Mechanisms

**DOI:** 10.1101/2022.06.24.497461

**Authors:** Jordan T. Bateman, Erica S. Levitt

## Abstract

Opioids depress breathing by inhibition of inter-connected respiratory nuclei in the pons and medulla. Mu opioid receptor (MOR) agonists directly hyperpolarize a population of neurons in the dorsolateral pons, particularly the Kölliker-Fuse (KF) nucleus, that are key mediators of opioid-induced respiratory depression. However, the projection target and synaptic connections of MOR-expressing KF neurons is unknown. Here, we used retrograde labeling and brain slice electrophysiology to determine that MOR-expressing KF neurons project to respiratory nuclei in the ventrolateral medulla, including the pre-Bötzinger complex (preBötC) and rostral ventral respiratory group (rVRG). These medullary projecting, MOR-expressing dorsolateral pontine neurons express FoxP2 and are distinct from calcitonin gene-related peptide-expressing lateral parabrachial neurons. Furthermore, dorsolateral pontine neurons release glutamate onto excitatory preBötC and rVRG neurons via monosynaptic projections, which is inhibited by presynaptic opioid receptors. The excitatory preBötC and rVRG neurons receiving MOR-sensitive glutamatergic synaptic input from the dorsolateral pons are themselves hyperpolarized by opioids. Thus, opioids can synergistically inhibit this excitatory pontomedullary respiratory circuit by three distinct mechanisms—somatodendritic MORs on dorsolateral pontine and ventrolateral medullary neurons and presynaptic MORs on dorsolateral pontine neuron terminals in the ventrolateral medulla—all of which could contribute to opioid-induced respiratory depression.

## INTRODUCTION

With the prevalence of opioid overdose on the rise (Wilson *et al*., 2020; Mattson *et al*., 2021), understanding the mechanisms of opioid-induced respiratory depression is of particular importance to aid in development of countermeasures and/or analgesics that do not affect breathing. Opioids, due to activation of the mu opioid receptor (MOR) (Dahan *et al*., 2001), depress breathing by inhibiting inter-connected respiratory nuclei in the pons and medulla (Bateman *et al*., 2021; Ramirez *et al*., 2021). Despite significant progress, detailed mechanisms by which this occurs remain elusive, especially for the dorsolateral pons. We sought to identify mechanistic insight concerning how opioids inhibit pontomedullary respiratory neurocircuitry that gives rise to opioid-induced respiratory depression.

Respiration is generated and controlled by an interconnected pontomedullary network in the brainstem (Del Negro *et al*., 2018). The Kölliker-Fuse (KF) nucleus and adjacent lateral parabrachial area (LPB) of the dorsolateral pons are critical to the neural control of breathing (Lumsden, 1923; Fung & St John, 1995; Dutschmann & Herbert, 2006; Smith *et al*., 2007). The KF/LPB is composed of a heterogeneous population of respiratory neurons that impact respiratory rate and pattern (Chamberlin & Saper, 1994; Navarrete-Opazo *et al*., 2020; Saunders & Levitt, 2020) via excitatory projections to respiratory nuclei in the ventrolateral medulla, including, but not limited to the Bötzinger Complex (BötC), preBötzinger Complex (preBötC), and rostral ventral respiratory group (rVRG) (Song *et al*., 2012; Yokota *et al*., 2015; Geerling *et al*., 2017; Yang *et al*., 2020). The preBötC generates inspiratory rhythm (Smith *et al*., 1991), which is relayed to inspiratory premotor neurons in the rVRG. The BötC contains mostly inhibitory neurons that fire during expiration and is a major source of inhibition within the network (Schreihofer *et al*., 1999; Ezure *et al*., 2003). The dynamic interplay between the KF/LPB and the BötC, preBötC, and rVRG is essential for optimized respiratory output (Dutschmann & Dick, 2012; Smith *et al*., 2007). Unfortunately, all of these respiratory nuclei express MORs leading to inhibition of the control of breathing network via multiple potential sites and mechanisms (Gray *et al*., 1999; Lonergan *et al*., 2003; Montandon *et al*., 2011; Levitt *et al*., 2015; Cinelli *et al*., 2020).

The two respiratory nuclei considered most critical for opioid-induced respiratory depression are the KF/LPB of the dorsolateral pons and the preBötC of the ventrolateral medulla (Bachmutsky *et al*., 2020; Varga *et al*., 2020). The dorsolateral pontine KF/LPB is considered a key contributor of opioid-induced respiratory depression because: 1) deletion of MORs from the KF/LPB attenuates morphine-induced respiratory depression (Bachmutsky *et al*., 2020; Varga *et al*., 2020; Liu *et al*., 2021), 2) opioids injected into the KF/LPB reduce respiratory rate (Prkic *et al*., 2012; Levitt *et al*., 2015; Miller *et al*., 2017; Liu *et al*., 2021), 3) blockade of KF/LPB opioid receptors rescues fentanyl-induced apnea (Saunders & Levitt, 2020), and 4) chemogenetic inhibition of MOR-expressing LPB neurons induces respiratory depression (Liu *et al*., 2021). Yet, mechanisms by which the opioid inhibition of dorsolateral pontine neurons alter neurotransmission in the respiratory circuitry and causes suppression of breathing are unknown.

MORs inhibit neurotransmission by hyperpolarizing neurons and/or inhibiting presynaptic neurotransmitter release (Jiang & North, 1992; Chahl, 1996; Zamponi & Snutch, 1998; Al-Hasani & Bruchas, 2011). In the preBötC, presynaptic MORs inhibit synaptic transmission (Ballanyi *et al*., 2010; Wei & Ramirez, 2019; Baertsch *et al*., 2021) and are expressed more abundantly than somatodendritic MORs (Lonergan *et al*., 2003). These presynaptic MORs in the preBötC are poised to play a major role in the mechanism of opioid suppression of breathing within the inspiratory rhythm generating area, but the specific origins of MOR-expressing synaptic projections remains unknown. Here we tested the hypothesis that they are coming from the dorsolateral pons.

Opioids hyperpolarize a subset of KF neurons (Levitt *et al*., 2015), whose neurochemical identity and possible projection targets are unknown. Glutamatergic KF neurons project to the ventrolateral medulla (Song *et al*., 2012; Yokota *et al*., 2015; Geerling *et al*., 2017) and, if inhibited by opioids—either by somatodendritic activation of GIRK channels and/or presynaptic inhibition of neurotransmitter release—could depress breathing. Therefore, we hypothesized that MOR-expressing KF neurons project to and form excitatory synapses onto respiratory controlling neurons in the ventrolateral medulla (i.e. the preBötC and rVRG), and that this excitatory synapse is inhibited by presynaptic MORs on KF terminals. The results show that this excitatory pontomedullary respiratory circuit is robustly inhibited by opioids by three different mechanisms, involving presynaptic and postsynaptic opioid receptors in the dorsolateral pons and the ventrolateral medulla, revealing convergent mechanisms by which opioids can depress breathing.

## RESULTS

### Oprm1 expression in dorsolateral pontine neurons

To visualize Oprm1-expressing dorsolateral pontine neurons, Oprm1^Cre/Cre^ mice (Baertsch *et al*., 2021; Liu *et al*., 2021) were crossed with Ai9 tdTomato Cre-reporter mice to generate Ai9^tdt+^::oprm1^Cre/+^ mice (hereby referred to as Oprm1^Cre/tdT^ mice) that express tdTomato in neurons that also express MORs at any point during development. Oprm1-expressing neurons and neurites were identified in the dorsolateral pons, specifically in the lateral parabrachial area and KF (n=3; Figure 1A-D).

**Figure 1.**
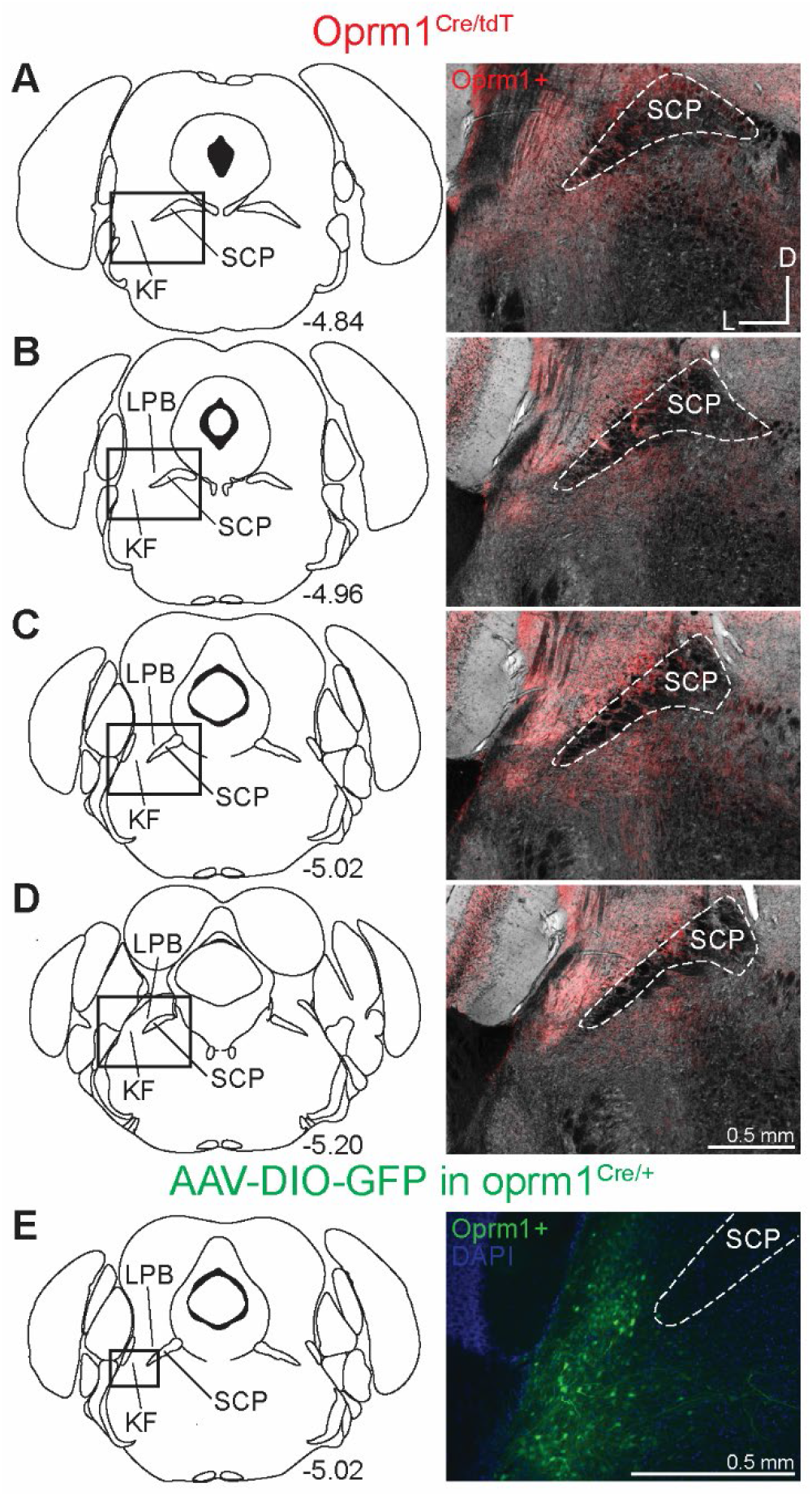
Dorsolateral pontine neurons express MORs. **(A-D)** Representative images of tdTomato expression, as an indicator of MOR expression, in coronal dorsolateral pontine slices from Oprm1^Cre/tdT^ mice (n = 3) across the rostral to caudal KF/LPB axis. Fluorescent tdTomato image is overlaid onto brightfield image to show landmarks. **(E)** Representative image following injection of virus encoding Cre-dependent GFP expression into KF/LPB to label MOR+ neurons in adult Oprm1^Cre/+^ mice (n=5). The approximate levels caudal to bregma are to the right of each schematic. The scale bar in (D) applies to images (A-D). Kölliker-Fuse (KF), lateral parabrachial (LPB), superior cerebellar peduncle (SCP).

To selectively label neurons that express MORs during adulthood, a virus encoding Cre-dependent GFP expression (AAV-DIO-GFP) was injected into the dorsolateral pons of Oprm1^Cre/+^ 2-4-month-old mice (n = 5). Oprm1-expressing neurons were again identified in the lateral parabrachial and KF areas (Figure 1E). Neuronal cell bodies were more apparent in these images since MOR-expressing afferents into the dorsolateral pons were not labeled by this approach. These results are consistent with previous studies showing that MORs are expressed in LPB (Huang *et al*., 2021; Liu *et al*., 2021) and KF (Levitt *et al*., 2015; Varga *et al*., 2020).

### Oprm1+ KF neurons project to respiratory nuclei in the ventrolateral medulla

We hypothesized that Oprm1+ KF neurons project to respiratory controlling nuclei in the ventrolateral medulla, especially the preBötC and rVRG. To determine this, retrograde virus encoding Cre-dependent expression of GFP (retrograde AAV-hSyn-DIO-eGFP) was unilaterally injected into the preBötC or the rVRG of Oprm1^Cre/+^ mice (Figure 2). As a control, anterograde virus encoding mCherry (AAV2-hSyn-mCherry) was co-injected to mark the injection site. The intensity of mCherry expression was measured throughout the rostral-caudal axis of the ventrolateral medulla to quantify the extent of injection spread in accordance with medullary anatomical markers (Figure 2B and C). In addition, immunolabeling for the neurokinin 1 receptor (NK1R) was used as a marker of the preBötC (Gray *et al*., 1999; Montandon *et al*., 2011) and to identify the nucleus ambiguous (NA), which was especially useful for the compact section of the NA in the preBötC region (Figure 2B (bottom)). Injection sites were categorized based on the location of peak mCherry expression intensity (Figure 2C & Figure 2 – Figure supplement 1).

**Figure 2.**
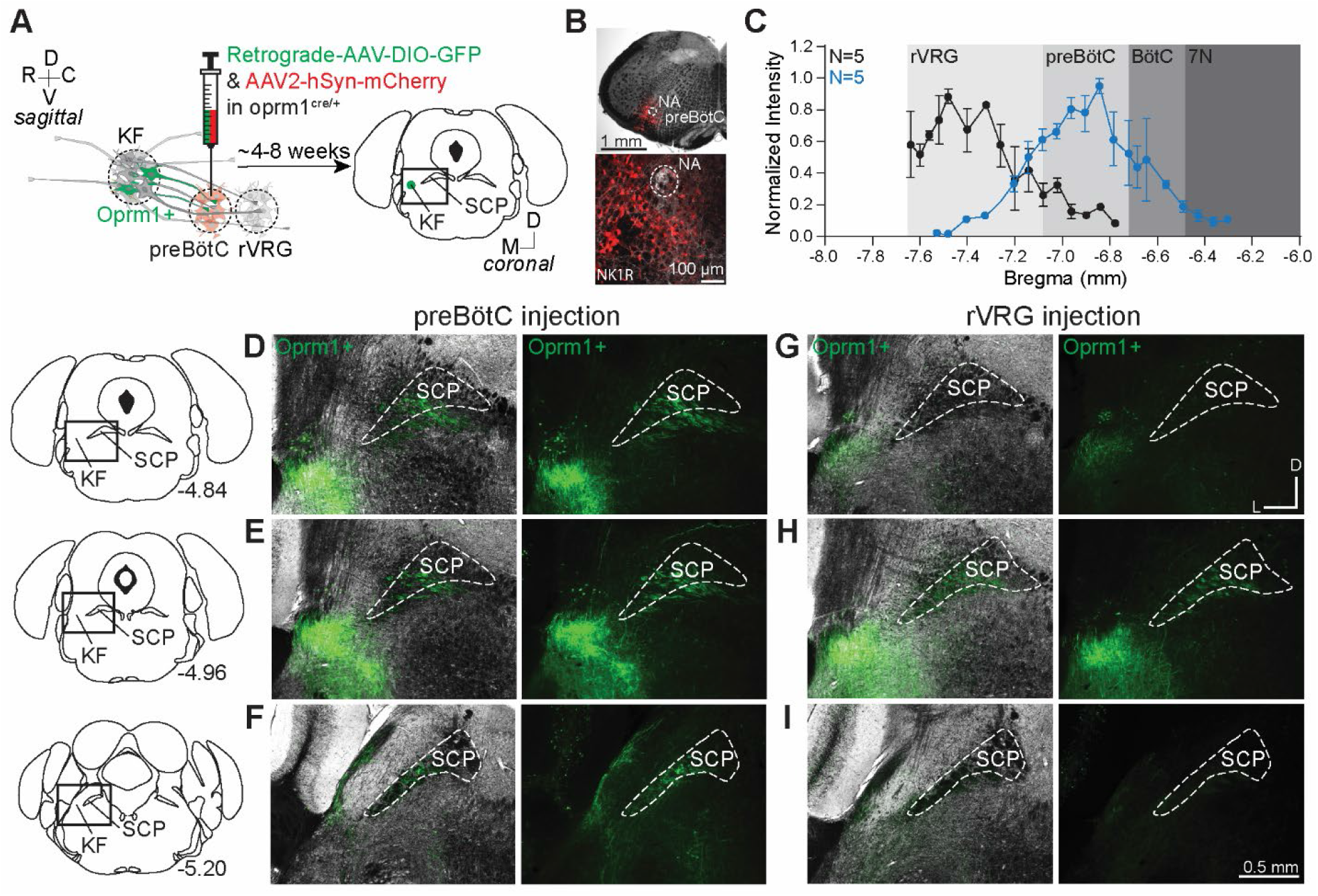
Oprm1+ KF neurons project to the preBötC and rVRG. **(A**) Schematic illustrating the approach to retrogradely label Oprm1+ KF neurons projecting to the preBötC or rVRG. **(B)** Images of coronal slices from the medulla with a control injection of AAV2-hSyn-mCherry into the preBötC of an Oprm1^Cre/+^ mouse to mark the injection site. Immunohistochemistry for the neurokinin 1 receptor (NK1R) was used as a marker for the preBötC and the nucleus ambiguous (NA). **(C)** Quantification of normalized AAV2-hSyn-mCherry fluorescence intensity along the rostral to caudal axis in the ventrolateral medulla of preBötC (n=5) and rVRG (n=5). Anatomical level relative to Bregma is indicated on the x-axis. **(D-I)** Representative images of GFP expression, as an indicator of retrograde labeled Oprm1-expressing neurons, following injections into the preBötC (D-F) or the rVRG (G-I) across three levels of the dorsolateral pons. The bregma level is indicated on the schematics. The scale bar in (I) applies to all images (D-I).

Oprm1+ dorsolateral pontine neurons were retrogradely labeled from both preBötC and rVRG (Figure 2 D-I). Interestingly, Oprm1+ neurons that project to the preBötC (Figure 2D-F; n = 5) and the rVRG (Figure 2G-I; n=5) were mostly localized to the rostral and mid-rostral KF, and nearly absent in the caudal KF and lateral parabrachial area (Figure 2F & 2I). The majority of the retrogradely labeled dorsolateral pontine neurons were ipsilateral to the injection site, with very few or no contralateral neurons expressing GFP. Injections in three mice were located rostrally from the preBötC with the peak of mCherry expression in the BötC (Figure 2 – Figure supplement 1). In contrast to preBötC and rVRG projections, qualitatively fewer Oprm1+ KF neurons projected to the BötC (Figure 2 – Figure supplement 1E-G).

### Presynaptic opioid receptors inhibit glutamate release from KF terminals onto excitatory medullary neurons

Using anterograde labeling of Oprm1+ KF neurons (AAV2-DIO-ChR2-GFP into the KF of Oprm1^Cre/+^ mice), we found abundant Oprm1+ projections in the preBötC area (Figure 3A; n=4). Given this and knowledge that the KF neurons projecting to the ventrolateral medulla are glutamatergic (Geerling *et al*., 2017) and express MORs (Figure 2), we hypothesized that opioids inhibit glutamate release from KF terminals onto respiratory neurons in the ventrolateral medulla, particularly the preBötC and rVRG. To test this hypothesis, we unilaterally injected a virus encoding channelrhodopsin2 (AAV2-hSyn-hChR2(H134R)-EYFP-WPRE-PA) into the KF of vglut2^Cre/tdT^ mice (Figure 3B & 3C). We made whole-cell voltage clamp recordings from tdTomato-expressing, excitatory vglut2-expressing preBötC and rVRG neurons contained in acute brain slices (Figure 3D). We chose to target vglut2-expressing neurons since 1) this contains the population of inspiratory rhythm-generating preBötC neurons (Wallén-Mackenzie *et al*., 2006; Gray *et al*., 2010; Cui *et al*., 2016) and inspiratory premotor rVRG neurons, 2) KF neurons project to excitatory, more so than inhibitory, preBötC neurons (Yang *et al*., 2020), and 3) deletion of MORs from vglut2 neurons eliminates opioid-induced depression of respiratory output in medullary slices (Sun *et al*., 2019; Bachmutsky *et al*., 2020). Optogenetic stimulation of KF terminals drove pharmacologically-isolated excitatory postsynaptic currents (oEPSCs) in excitatory preBötC and rVRG neurons (Figure 3E & 3H) that were blocked by the AMPA-type glutamate receptor antagonist DNQX (6,7-dinitroquinoxaline-2,3-dione; 10 μM; Figure 3K & 3M, n=11). Additionally, KF synapses onto medullary respiratory neurons are monosynaptic because oEPSCs were eliminated by tetrodotoxin (TTX; 1μM) yet restored by subsequent application of 4-aminopyridine (4AP; 100 μM) (Figure 3K & 3L; n=7). Thus, KF neurons send monosynaptic, glutamatergic projections to excitatory ventrolateral medullary neurons.

**Figure 3.**
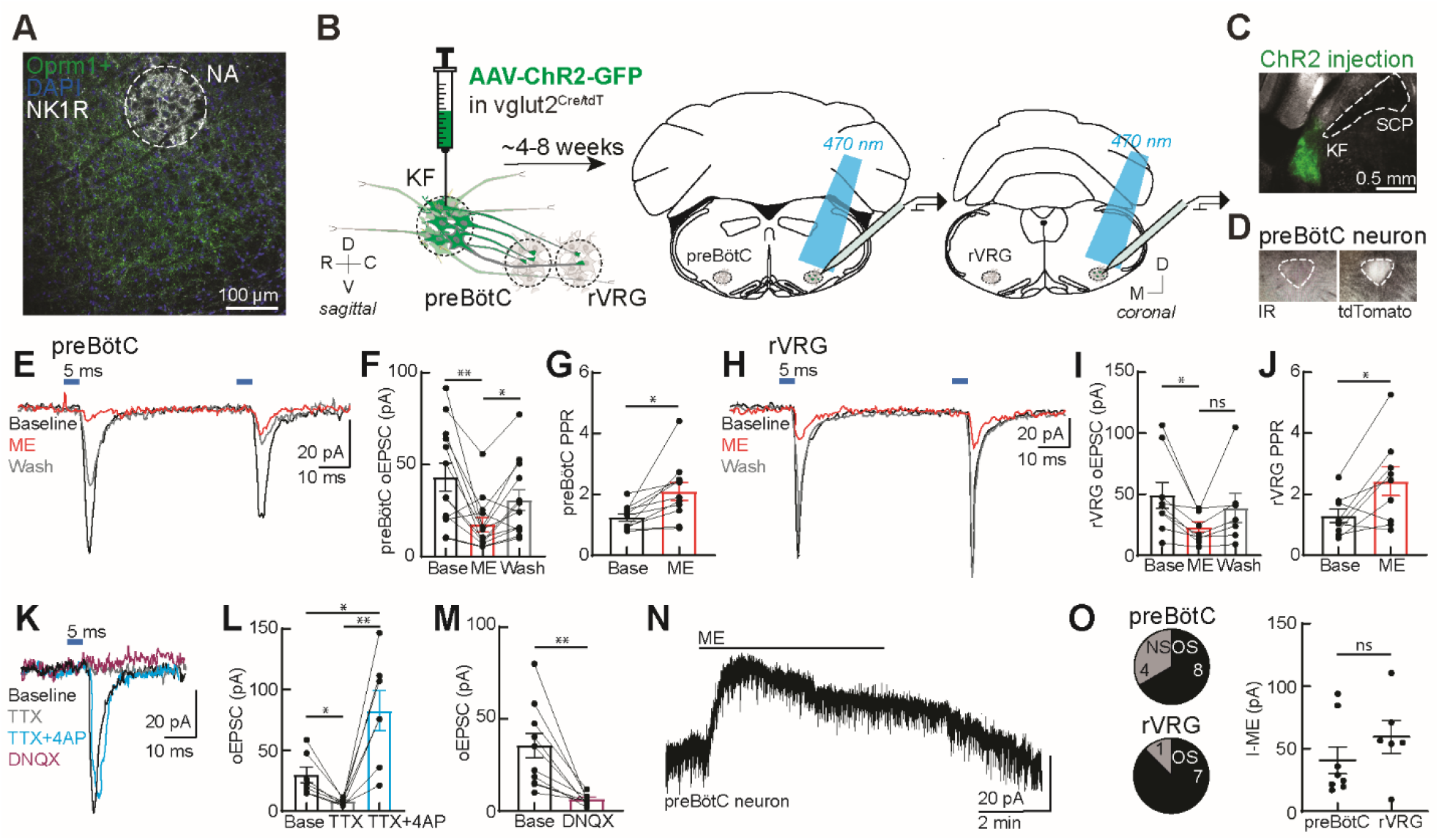
Presynaptic opioid receptors inhibit glutamate release from KF terminals onto excitatory preBötC and rVRG neurons. **(A)** Representative image of Oprm1+ projections from dorsolateral pons to the preBötC labeled from injection of AAV2-DIO-ChR2-GFP into the KF of Oprm1^Cre/+^ mice (n=4). Immunolabeling for NK1R was used as an anatomical marker of the NA. **(B)** Schematic of approach to optogenetically stimulate KF terminals and drive oEPSCs in excitatory preBötC and rVRG neurons in an acute brain slice. **(C)** Representative image of ChR2-GFP expression in the KF (injection area) of vglut2^Cre/tdT^ mouse. **(D)** tdTomato-expressing, excitatory vglut2-expressing preBötC (or rVRG) neurons were identified in acute brain slices. **(E)** Recording of pairs of oEPSCs (5 ms stimulation, 50 ms inter-stimulus interval) from an excitatory preBötC neuron in an acute brain slice at baseline (black), during perfusion of ME (3 μM) (red), and after wash (gray). **(F)** ME decreased oEPSC amplitude in preBötC neurons (n = 13; **p=0.007, *p=0.013 by one-way ANOVA and Tukey’s post-test). **(G)** ME increased the paired-pulse ratio (P2/P1) in preBötC neurons (n=11; *p = 0.001 paired t-test). **(H)** Recording of pairs of oEPSCs (5 ms stimulation, 50 ms inter-stimulus interval) from an excitatory rVRG neuron in an acute brain slice at baseline (black), during perfusion of ME (3 μM) (red), and after wash (gray). **(I)** ME decreased oEPSC amplitude in rVRG neurons (n = 9; *p=0.027 by one-way ANOVA and Tukey’s post-test). **(J)** ME increased the paired-pulse ratio (P2/P1) in rVRG neurons (n=9; *p=0.043 by paired t-test). **(K)** Recording of oEPSCs from an excitatory preBötC neuron in an acute brain slice at baseline (black), during perfusion of TTX (1 μM) (gray), during perfusion of TTX (1 μM) + 4AP (100 μM) (cyan), and during perfusion of DNQX (10 μM) (purple). **(L)** KF synapses onto medullary respiratory neurons are monosynaptic. TTX blocked oEPSCs (n = 7; *p=0.0207 by one-way ANOVA and Tukey’s post-test), which were restored by perfusion of TTX + 4AP (n = 7; **p=0.0086, *p=0.042 by one-way ANOVA and Tukey’s post-test). **(M)** KF synapses onto medullary respiratory neurons are glutamatergic. AMPA-type glutamate receptor antagonist DNQX (10 μM) blocked oEPSCs (n=11; **p=0.001 paired t-test). **(N)** Recording from a preBötC neuron in an acute brain slice. ME (3 μM) induced an outward current. **(O)** ME-mediated outward currents were observed in 8 of 12 preBötC neurons and 6 of 7 rVRG neurons. OS=opioid-sensitive, NS=non-opioid-sensitive. There was no difference in the amplitude of ME-mediated currents in OS preBötC (n=8) and rVRG neurons (n=6) (ns, p=0.294; unpaired t-test). For all graphs, bar/line and error represent mean ± SEM. Individual data points are from individual neurons.

To determine whether opioids inhibit glutamate release from KF terminals onto medullary respiratory neurons, pairs of oEPSCs (50 ms inter-stimulus interval) were recorded from excitatory preBötC and rVRG neurons, and the endogenous opioid agonist [Met^5^]enkephalin (ME) was applied to the perfusion solution. ME (3 μM) decreased the oEPSC amplitude in preBötC neurons (Figure 3E & 3F; n=13) and in rVRG neurons (Figure 3H & 3I; n=9), which reversed when ME was washed from the slice (Figure 3F & 3I). The opioid-sensitivity of glutamate release from KF terminals was fairly homogeneous, considering that oEPSCs were inhibited by ME by at least 30% in nearly all preBötC neurons (11 of 13 neurons) and all rVRG neurons. In addition, ME increased the paired-pulse ratio (PPR) in both preBötC (Figure 3G; n=11) and rVRG neurons (Figure 3G; n=9), indicating inhibition of glutamate release by presynaptic MORs. Thus, presynaptic opioid receptors inhibit glutamate release from KF terminals onto excitatory preBötC and rVRG neurons.

We were also able to determine whether the excitatory preBötC or rVRG neuron that received opioid-sensitive glutamatergic synaptic input from the KF was itself hyperpolarized by opioids by monitoring the holding current. ME (3 μM) induced an outward current (Figure 3N) in 68% of preBötC neurons (8 of 12 neurons) and 88% of rVRG neurons (7 of 8 neurons) (Figure 3O). There was no difference in the amplitude of the ME-mediated current in preBötC and rVRG neurons (Figure 3O). Thus, a majority of excitatory preBötC and rVRG neurons that receive opioid-sensitive glutamatergic synapses from KF neurons are themselves hyperpolarized by opioids, indicating both pre and postsynaptic suppression of this excitatory synapse by opioids.

### Opioids hyperpolarize medullary-projecting KF neurons

Opioids hyperpolarize a subpopulation of KF neurons by activating G protein-coupled inwardly rectifying potassium (GIRK) channels (Levitt *et al*., 2015). Given that Oprm1+ KF neurons that project to the ventrolateral medulla express functional MORs on presynaptic terminals (Figures 2 and 3), we wanted to determine whether these neurons also express functional somatodendritic MORs leading to hyperpolarization. We recorded from KF neurons retrogradely labeled with FluoSpheres (580/605) that were unilaterally injected into the preBötC or rVRG of wild-type mice (Figure 4A). FluoSpheres were chosen over viral retrograde tracers for these experiments because they are highly visible in acute brain slices and do not spread as far in the injection area (Figure 4A), genetically alter neurons, require fluorescent amplification, or take long to express (2 days vs 4 weeks). Furthermore, FluoSpheres will label KF neurons regardless of Oprm1 expression status, enabling us to determine the projection pattern of both Oprm1+ and Oprm1-neurons. Whole-cell voltage-clamp recordings were made from fluorescent KF neurons contained in acute brain slices (Figure 4B). The presence of an ME-mediated outward current identified KF neurons that express functional MORs and were opioid sensitive (OS) (Figure 4C), as compared to neurons that lacked an ME-mediated outward current (non-sensitive; NS) (Figure 4D). ME induced an outward current in 59% (13 of 22 neurons) of KF neurons that project to the preBötC (Figure 4E) and 67% (12 of 18 neurons) of KF neurons that project to the rVRG (Figure 4F). The average amplitude of the ME-mediated current was not different between KF neurons that project to preBötC (n = 13) or rVRG (n = 12) (p=0.8250; unpaired *t* test). Thus, both opioid sensitive and non-sensitive KF neurons project to preBötC and rVRG.

**Figure 4.**
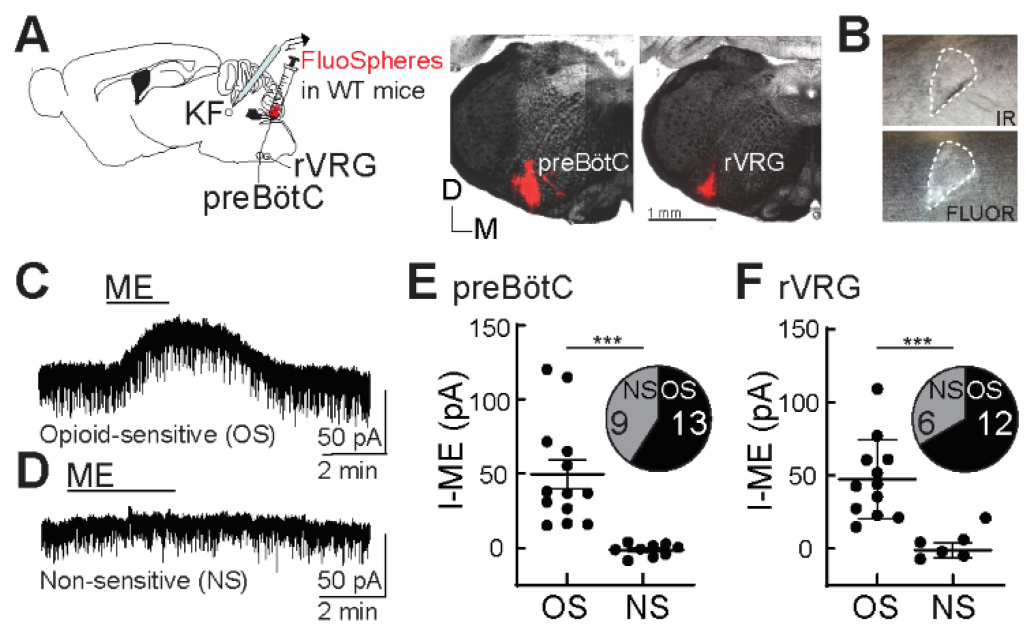
Opioids hyperpolarize KF neurons that project to the preBötC and rVRG. **(A)** Schematic (left) of approach to retrogradely label KF neurons that project to the preBötC or rVRG with FluoSpheres in wild-type mice. Images (right) of FluoSpheres in the injection area (preBötC or rVRG). The scale bar applies to both injection images. **(B)** A KF neuron retrogradely labeled by FluoSpheres shown with IR-Dodt and epifluorescent (FLUOR) illumination. **(C&D)** Whole-cell voltage-clamp recordings from opioid-sensitive (E, “OS”) and non-opioid-sensitive (F, “NS”) retrogradely-labeled KF neurons. ME (1 µM) induced an outward current in the opioid-sensitive (OS) neuron (E), but not the non-opioid-sensitive (NS) neuron (F). **(E&F)** Quantification of the amplitude of the ME-mediated current (I-ME (pA)) in opioid-sensitive and non-opioid-sensitive KF neurons that project to the preBötC (H; n=22; ***p=0.0005; unpaired t-test) or the rVRG (I; n=18; ***p=0.0007; unpaired t-test). ME induced an outward current in 13 of 22 KF neurons that project to the preBötC and 12 of 18 KF neurons that project to the rVRG. Individual data points are from individual neurons in separate slices. Line and error are mean ± SEM.

Given the potentially lesser degree of projections from Oprm1+ KF neurons to the BötC (Figure 4 – Figure supplement 2) and the ability to retrogradely label Oprm1 negative neurons with FluoSpheres, we also injected FluoSpheres into the BötC (n = 11) to test the hypothesis that Oprm1 negative KF neurons project to the BötC (Figure 4 – Figure supplement 2). We made whole-cell voltage-clamp recordings from fluorescent KF neurons and found that ME induced an outward current in only 36% (4 of 11 neurons) of KF neurons that project to the BötC (Figure 4 – Figure supplement 2C). Thus, a lower proportion of opioid sensitive neurons project to BötC, compared to preBötC and rVRG.

### Distribution of Oprm1+ and Oprm1-dorsolateral pontine neurons projecting to the ventrolateral medulla

To further examine the distribution of Oprm1+ and Oprm1-dorsolateral pontine neurons projecting to the ventrolateral medulla, retrograde AAV-hSyn-DIO-eGFP and retrograde AAV-hSyn-mCherry were unilaterally injected into the preBötC and rVRG of Oprm1^Cre/+^ mice (Figure 5A). Using this approach, projection neurons that express Oprm1 will express GFP and mCherry, whereas projection neurons that do not express Oprm1 will only express mCherry (Figure 5B). The number of mCherry and/or GFP expressing neurons was evaluated in rostral (∼bregma level -4.84 mm), mid-rostral (∼bregma level -4.96 mm), and caudal (∼bregma level - 5.20 mm) sections of the dorsolateral pons (n=4 mice, 3 slices per region per mouse). There were significantly more retrograde labeled neurons in rostral and mid-rostral slices, regardless of Oprm1 expression status (Figure 5D). Consistent with previous observations (Figure 2), retrograde labeled Oprm1+ neurons were mostly localized to the rostral and mid-rostral slices, and not in caudal slices or lateral parabrachial area (Figure 5C&E; Figure 5 – Figure supplement 3). The percentage of retrograde labeled neurons that were Oprm1+ (co-labeled with mCherry and GFP) in rostral slices (56%) and mid-rostral slices (47%) was higher than in caudal slices (15%) (Figure 5C). Taken together, Oprm1+ and Oprm1-KF neurons that project to respiratory nuclei in the ventrolateral medulla are distributed to the rostral and mid-rostral regions of the KF of mice.

**Figure 5.**
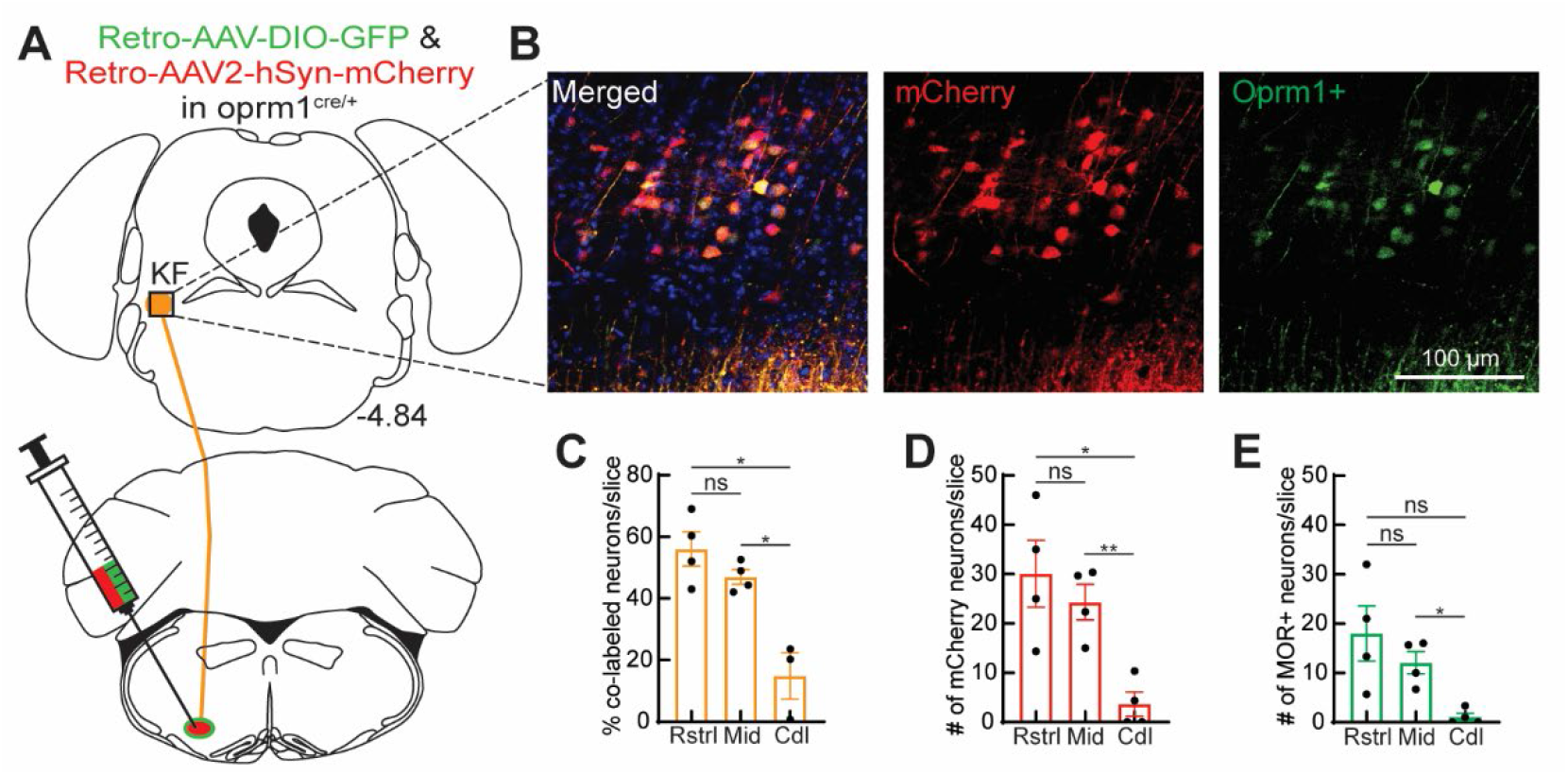
Oprm1+ and Opmr1-dorsolateral pontine neurons project to the ventrolateral medulla. **(A)** Schematic of approach injecting retrograde virus encoding Cre-dependent GFP expression and a retrograde virus encoding mCherry expression into the ventrolateral medulla of Oprm1^Cre/+^ mice to label Oprmr1+ and Oprm1-dorsolateral pontine neurons that project to these respiratory nuclei. **(B)** Representative images of mCherry expression (retrogradely labels neurons regardless of Oprm1 expression) and GFP expression (retrogradely labels Oprm1+ neurons) in a rostral dorsolateral pontine slice (bregma -4.84 mm). **(C)** Summary of percentage of retrograde-labeled neurons that were Oprm1+ (co-labeled with mCherry and GFP) in rostral (Rstrl, bregma -4.84 mm), mid-rostral (Mid, bregma -4.96 mm), and caudal (Cdl, bregma -5.2 mm) slices. **(D&E)** Summary of the average number of mCherry-expressing (D) or GFP-expressing MOR+ (E) dorsolateral pontine neurons per slice in rostral, mid-rostral, and caudal slices. Bar and error are mean ± SEM. Individual data points are from individual mice. N=4 mice, 3 slices per region per mouse. *p<0.05, **p<0.01, ns=p>0.05 by one-way ANOVA and Tukey’s post-test.

### Oprm1+, medullary-projecting KF neurons express FoxP2, but not CGRP

Rostral glutamatergic KF neurons express FoxP2 (Forkhead box protein P2) (Geerling *et al*., 2017; Karthik *et al*., 2022), whereas MOR-expressing glutamatergic neurons in the lateral parabrachial area that project to the forebrain express Calca, a gene that encodes the neuropeptide calcitonin gene-related peptide (CGRP) (Huang *et al*., 2021). Considering this, we performed immunohistochemistry for FoxP2 and CGRP on Oprm1+ KF neurons projecting to the ventrolateral medulla. Oprm1+, medullary-projecting KF neurons expressed FoxP2 (n=3; Figure 6), but not CGRP (n=3; Figure 7). Although CGRP expression was absent from the rostral KF and medullary projecting Oprm1+ neurons and neurites, it was robust in lateral parabrachial neurons and their axon fiber projections (Figure 7C).

**Figure 6.**
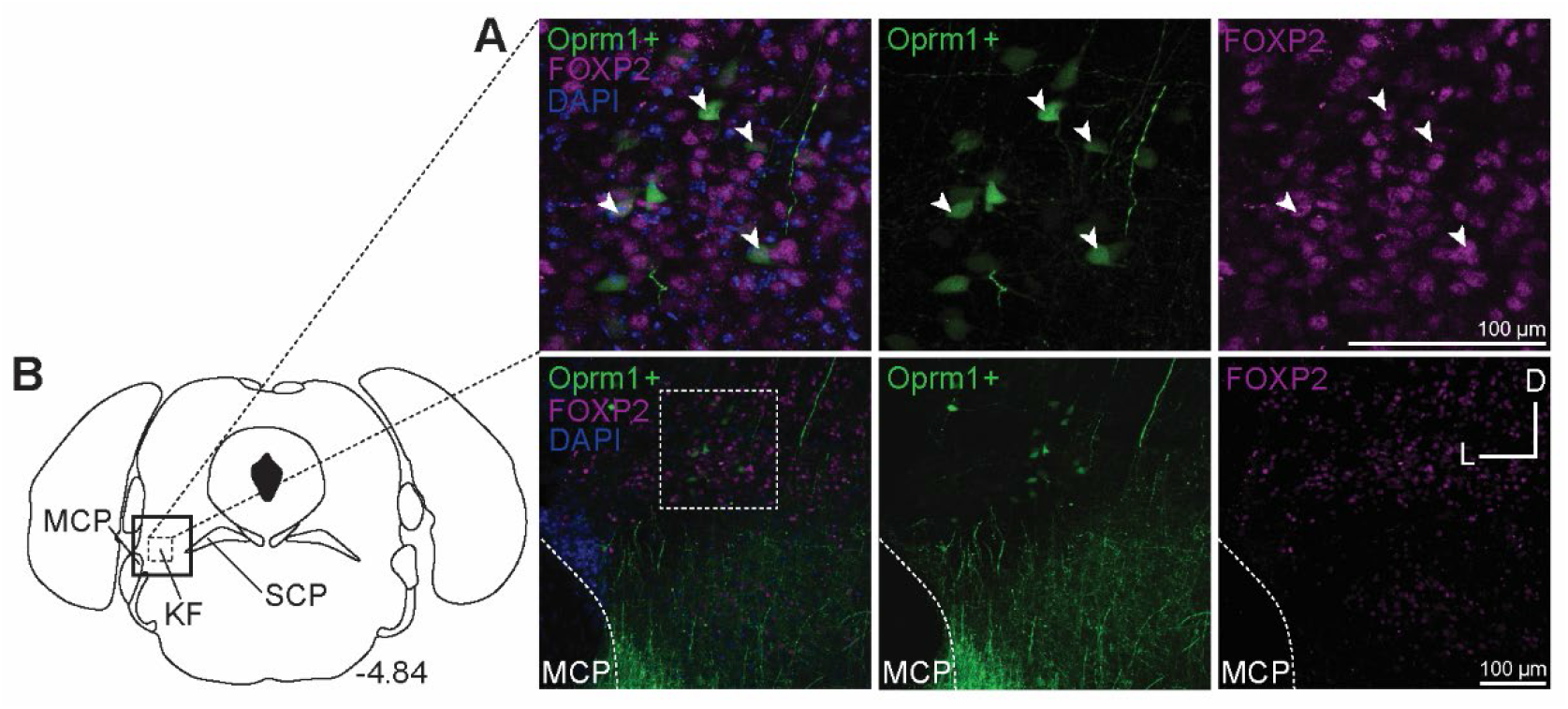
Oprm1+, medullary projecting KF neurons express FoxP2. Oprm1+ neurons that project to the ventrolateral medulla were retrogradely labeled by injection of retrograde AAV-DIO-GFP. Immunohistochemistry was used to label FoxP2. **(A-B)** In rostral slices, FoxP2 is expressed in Oprm1+ KF neurons that project to the ventrolateral medulla. The approximate bregma level is to the right of the schematic. The scale bars apply to all images in each row. FoxP2 (Forkhead box protein P2), SCP (superior cerebellar peduncle), and MCP (medial cerebellar peduncle).

**Figure 7.**
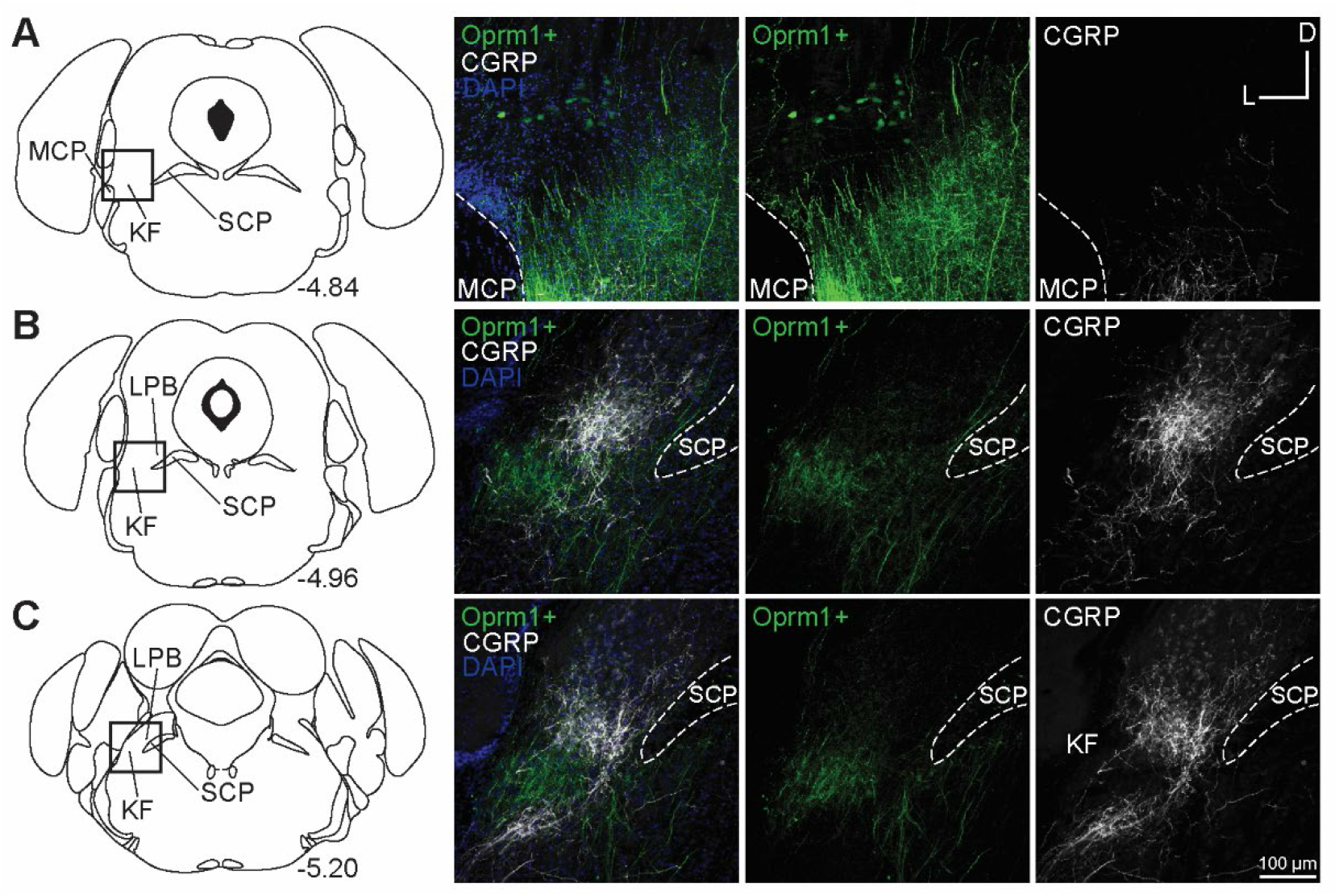
Oprm1+, medullary projecting KF neurons do not express CGRP. Oprm1+ neurons that project to the ventrolateral medulla were retrogradely labeled by injection of retrograde AAV-DIO-GFP. Immunohistochemistry was used to label CGRP. **(A)** CGRP is absent from rostral KF and Oprm1+ KF neurons that project to the ventrolateral medulla (Oprm1+). **(B & C)** CGRP marks LPB neurons and their axon fiber projections, but is absent from retrograde labeled Oprm1+ axon fiber projections in mid-rostral (B) and caudal (C) slices. The approximate bregma levels are to the right of each schematic. The scale bar in (C) applies to all images. CGRP (calcitonin gene-related peptide), SCP (superior cerebellar peduncle), and MCP (medial cerebellar peduncle).

## DISCUSSION

Opioid suppression of breathing could occur via multiple mechanisms and at multiple sites in the pontomedullary respiratory network. Here, we show that opioids inhibit an excitatory pontomedullary respiratory circuit via three mechanisms 1) postsynaptic MOR-mediated hyperpolarization of KF neurons that project to the ventrolateral medulla, 2) presynaptic MOR-mediated inhibition of glutamate release from dorsolateral pontine terminals onto excitatory preBötC and rVRG neurons, and 3) postsynaptic MOR-mediated hyperpolarization of the preBötC and rVRG neurons that receive pontine glutamatergic input. We targeted the excitatory vglut2-expressing neurons in the ventrolateral medulla because they contain the populations of inspiratory rhythm-generating preBötC neurons (Wallén-Mackenzie *et al*., 2006; Gray *et al*., 2010; Cui *et al*., 2016) and inspiratory premotor rVRG neurons, and MOR deletion from vglut2 neurons prevents opioid-induced respiratory depression in medullary slices (Sun *et al*., 2019; Bachmutsky *et al*., 2020). Opioid inhibition of excitatory drive from KF onto these respiratory neuron populations is important for rhythm generation (preBötC) and respiratory pattern formation (rVRG). Thus, there are convergent mechanisms of opioid-induced respiratory suppression, including both presynaptic and postsynaptic opioid receptors in the dorsolateral pons and the ventrolateral medulla.

These mechanistic insights are parsimonious with previous studies showing that MORs in the dorsolateral pons (Prkic *et al*., 2012; Levitt *et al*., 2015; Miller *et al*., 2017; Bachmutsky *et al*., 2020; Saunders & Levitt, 2020; Varga *et al*., 2020; Liu *et al*., 2021) and the preBötC (Gray *et al*., 1999; Sun *et al*., 2019; Bachmutsky *et al*., 2020; Varga *et al*., 2020) are key mediators of opioid-induced respiratory depression. Genetic deletion or pharmacological blockade of different subsets of pre and postsynaptic MORs in these areas mostly resulted in partial attenuation of opioid-induced respiratory rate suppression, presumably due to redundancy from the subset(s) of MORs that weren’t deleted or blocked. Additional MORs outside of this dorsolateral pontine to preBötC circuit likely contribute to respiratory suppression, since deletion of MORs from both dorsolateral pons and preBötC did not eliminate morphine-induced respiratory suppression (Bachmutsky *et al*., 2020). Other potential contributors include the caudal medullary raphe nuclei, since antagonism of dorsolateral pons, ventrolateral medulla AND caudal medullary raphe did eliminate remifentanil-induced respiratory depression (Palkovic *et al*., 2022), but mechanisms remain unclear.

Significant attention has been given to the mechanisms of opioid suppression of inspiratory rhythm generation in the preBötC (Sun *et al*., 2019; Bachmutsky *et al*., 2020; Baertsch *et al*., 2021). Presynaptic opioid receptors in the preBötC inhibit synaptic transmission and have been postulated to disrupt preBötC neuron bursting (Ballanyi *et al*., 2010; Wei & Ramirez, 2019; Baertsch *et al*., 2021) by inhibition of excitatory neurotransmission that is dominant during bursts (Ashhad & Feldman, 2020), but the projection-specific location(s) of these presynaptic MORs is unknown. Our study has revealed a projection-specific presence of presynaptic MORs on glutamatergic terminals from dorsolateral pontine inputs to the preBötC. Although other MOR-expressing glutamatergic inputs are also likely contributors, including collaterals within the preBötC (Rekling *et al*., 2000), the role of these specific pontine inputs on opioid inhibition of respiratory rhythm generation is worthy of further investigation.

Only a subpopulation of preBötC neurons contain MORs (Bachmutsky *et al*., 2020; Baertsch *et al*., 2021; Kallurkar *et al*., 2022). Single cell sequencing analysis estimated that only ∼8% of all preBötC neurons (Bachmutsky *et al*., 2020) or 3 of 17 of Dbx1-expressing inspiratory preBötC neurons (Kallurkar *et al*., 2022) express MORs. In contrast, using opto-tagging of neurons in rhythmic horizontal medullary slices from MOR-Cre mice, ∼50% of respiratory modulated preBötC neurons were estimated to express MORs (Baertsch *et al*., 2021). We previously found that ∼42% of preBötC neurons (8 of 19 neurons) were hyperpolarized by MORs (Varga et al., 2020). The population of MOR-expressing preBötC neurons is heterogeneous, including nearly equal numbers of glutamatergic, GABAergic, and glycinergic neurons (Bachmutsky *et al*., 2020), Type-1 and Type-2 Dbx1-expressing inspiratory neurons (Kallurkar *et al*., 2022), and pre-inspiratory, inspiratory, expiratory, and tonic neurons (Baertsch *et al*., 2021). We found that MOR-expressing dorsolateral pontine glutamatergic inputs seem to preferentially synapse onto MOR-expressing excitatory preBötC neurons, since 68% of preBötC neurons (8 of 12 neurons) that received glutamatergic input from the dorsolateral pons were hyperpolarized by opioid (Figure 3O). This percentage is higher than even the highest estimate of MOR-expressing preBötC neurons (Baertsch *et al*., 2021), suggesting dorsolateral pontine neurons preferentially target MOR-expressing glutamatergic preBötC neurons, which are important mediators of inspiratory rhythm generation and opioid-induced respiratory depression in medullary slices (Sun *et al*., 2019; Bachmutsky *et al*., 2020). This observation is also consistent with anatomical tracing studies showing that KF neurons project to excitatory, more so than inhibitory, preBötC neurons (Yang *et al*., 2020).

Compared to the preBötC, much less is known about opioid effects in the rVRG. Here we showed that MOR-expressing KF neurons project strongly to the rVRG (Figure 2&4) and form glutamatergic synapses onto excitatory rVRG neurons, which are inhibited by presynaptic MORs (Figure 3). Most of the excitatory rVRG neurons that receive glutamatergic input from the dorsolateral pons are hyperpolarized by opioid agonist (Figure 3). These mechanisms could contribute to suppression of rate and amplitude of phrenic nerve bursting induced by opioids in the rVRG (Lonergan *et al*., 2003; Cinelli *et al*., 2020).

Unexpectedly, KF neurons were more likely inhibited by presynaptic vs. somatodendritic MORs. Most KF terminals expressed presynaptic MORs, since ME inhibited glutamate release onto nearly all preBötC neurons and all rVRG neurons (Figure 3). In contrast, postsynaptic (somatodendritic) MOR-mediated outward currents were only observed in about two-thirds of medullary projecting KF neurons (Figure 4). There are multiple possible reasons for this apparent heterogeneity. First, KF neurons may express MORs more abundantly on terminals than in the somatodendritic region. Second, KF neurons that did not have outward currents and were deemed not-sensitive to opioids may express MORs, but lack GIRK channels, the functional readout we used to assess opioid sensitivity. However, this seems unlikely since the percentage of retrograde labeled neurons that were Oprm1+ (56% in rostral, and 47% in mid-rostral slices; Figure 5) nearly matched the percentages of functionally identified opioid-sensitive KF neurons (59% of preBötC projecting and 67% of rVRG projecting neurons; Figure 4). The last and most interesting possibility is that opioid-sensitive glutamatergic KF neurons preferentially synapse onto excitatory medullary neurons, while opioid-non-sensitive KF neurons are GABAergic and/or synapse onto non-excitatory (i.e. inhibitory) medullary neurons.

The dorsolateral pons includes the lateral parabrachial area and the KF, both of which have been implicated in opioid-induced respiratory depression (Levitt *et al*., 2015; Prkic *et al*., 2012; Varga *et al*., 2020; Liu *et al*., 2021). Although effects of MORs in the lateral parabrachial and KF areas appear similar, mechanisms likely differ since the neuronal populations have different projection patterns (Geerling *et al*., 2017; Huang *et al*., 2021) and are involved in different behaviors besides breathing, especially the lateral parabrachial area (Campos *et al*., 2018; Chen *et al*., 2018); Liu *et al*., 2022). In addition, the distinction between KF and lateral parabrachial area is not clear cut, though recent description of transcription factor expression in the dorsolateral pons provides opportunity to improve this (Karthik *et al*., 2022). MORs in KF neurons could reduce respiratory rate by decreasing excitatory input to the preBötC, as shown here. In the lateral parabrachial area, MOR-expression highly overlaps with expression of CGRP (Huang *et al*., 2021). CGRP-expressing lateral parabrachial neurons project primarily to the forebrain (Huang *et al*., 2021), and can stimulate breathing following CO2-induced arousal (Kaur *et al*., 2017). We found that MOR-expressing dorsolateral pontine neurons that project to the ventrolateral medulla are distinct from CGRP-expressing neurons (Figure 6), although a small population of CGRP-expressing, ventrolateral medulla projecting neurons have been reported (Huang *et al*., 2021). Instead, the ventrolateral medulla projecting dorsolateral pontine neurons expressed FoxP2 (Figure 6), and may constitute the population of glutamatergic FoxP2 and Lmx1b neurons recently described in the rostral KF (Karthik *et al*., 2022).

In conclusion, our results show that opioids inhibit an excitatory pontomedullary respiratory circuit by 3 distinct mechanisms—somatodendritic MORs on dorsolateral pontine and ventrolateral medullary neurons and presynaptic MORs on glutamatergic dorsolateral pontine axon terminals in the ventrolateral medulla—all of which could contribute to the profound effects of opioids on breathing.

## METHODS

### Animals

All experiments were approved by the Institutional Animal Care and Use Committee at the University of Florida (protocol #09515) and were in agreement with the National Institutes of Health “Guide for the Care and Use of Laboratory Animals.” Homozygous Oprm1^Cre/Cre^ mice (Liu et al., 2021) (Jackson Labs Stock #035574, obtained from Dr. Richard Palmiter, University of Washington) were crossed with homozygous Ai9-tdTomato Cre reporter mice (Jackson Labs Stock #007909) to generate Oprm1^Cre/tdT^ mice. Homozygous vglut2-ires-Cre mice (Jackson Labs Stock #028863) were crossed with homozygous Ai9-tdTomato Cre reporter mice (Jackson Labs Stock #007909) to generate vglut2^Cre/tdT^ mice. Oprm1^Cre/+^, Oprm1^Cre/tdT^, vglut2^Cre/tdT^ and wild-type C57BL/6J mice (male and female, 1-3 months old) were used for all experiments. Mice were bred and maintained at the University of Florida animal facility. Mice were group-housed with littermates in standard sized plastic cages and kept on a 12 h light–dark cycle, with water and food available *ad libitum*.

### Stereotaxic injections

Mice were anesthetized with isoflurane (2-4% in 100% oxygen; Zoetis, Parsippany-Troy Hills, NJ) and placed in a stereotaxic alignment system (Kopf Instruments model 1900, Tujunga, CA). The dorsal skull was exposed and leveled horizontally in preparation for a small, unilateral craniotomy targeting either the KF (y = -5 mm, x = ± 1.7 mm, z = -3.9 mm from bregma), BötC (y = -6.6 mm and x = ± 1.3 mm from bregma, z = - 5.625 mm), preBötC (y = -6.9 mm and x = ± 1.3 mm from bregma, z = - 5.625 mm), or rVRG (y = -7.2 mm and x = ± 1.3 mm from bregma, z = - 5.625 mm). Virus (undiluted) or FluoSpheres (580/605, 0.04 µm, diluted to 20% in saline, Invitrogen) were loaded into freshly pulled glass micropipettes and injected using a Nanoject III pressure injector (Drummond Scientific Company, Broomall, PA, USA) at a rate of 10 nl every 20 seconds (100-200 nl total). Following the injection, the pipette was left in place for 10 minutes and slowly retracted. The wound was closed using Vetbond tissue adhesive (3M Animal Care Products, St Paul, MN, USA). Mice received meloxicam (5 mg kg^− 1^ in saline, s.c.) and were placed in a warmed recovery chamber until they were ambulating normally. Mice were used either 2-6 days (FluoSpheres) or 4-5 weeks (virus) later for electrophysiology, microscopy, or immunohistochemistry.

For retrograde labeling in Oprm1^Cre/+^ mice, a 1:1 mixture (100 nl total) of either retrograde AAV-hSyn-DIO-eGFP (Addgene) and AAV2-hSyn-mCherry (UNC vector core) (Figures 2, 6, & 7; Figure 2 – Figure supplement 1) or retrograde AAV-hSyn-DIO-eGFP (Addgene) and retrograde AAV-hSyn-mCherry (Addgene) (Figure 5) was injected into the BötC, preBötC and/or the rVRG. For labeling Oprm1+ KF neurons, AAV2-hSyn-DIO-EGFP (Addgene; 100 nl) (Figure 1E) or AAV2-hSyn-DIO-EGFP (Addgene; 100 nl) (Figure 3A) was injected into the KF of Oprm1^Cre/+^ mice. Vglut2^Cre/tdT^ mice received AAV2-hSyn-hChR2(H134R)-EYFP-WPRE-PA (Addgene; 100 nL) injections targeting the KF (Figure 3). Lastly, FluoSpheres (580/605, diameter: 0.04 µm, 20% in saline; 100 nL) were unilaterally injected into the BötC (Figure 4 – Figure supplement 2), preBötC or rVRG of wild type C57BL/6J mice (Figure 4).

The correct placement of injections into the either the KF, BötC, preBötC or rVRG was verified by anatomical landmarks, immunohistochemistry and fluorescence in free-floating coronal brain slices (40-100 µm) using a MultiZoom microscope (Nikon AZ100). The BötC, preBötC and rVRG are located bilaterally in a rostro-caudal column in the ventrolateral medulla, just ventral to the nucleus ambiguus. The BötC, preBötC and rVRG can be distinguished using the inferior olives, nucleus ambiguus, and nucleus tractus solitarius as medullary landmarks (Franklin & Paxinos, 2008; Varga *et al*., 2020). The KF is located bilaterally in the dorsolateral pons, just ventrolateral to the tip of the superior cerebellar peduncle and medial of the middle cerebellar peduncle (Varga *et al*., 2020; Karthik *et al*., 2022).

### Brain slice electrophysiology

Brain slice electrophysiology recordings were performed from KF neurons in acute brain slices from wild-type C57BL/6J mice or from vglut2-expressing preBötC and rVRG neurons in acute brain slices from vglut2^Cre/tdT^ mice injected with AAV2-hSyn-hChR2(H134R)-EYFP-WPRE-PA into the KF. Mice were anesthetized with isoflurane, decapitated, and the brain was removed and mounted in a vibratome chamber (VT 1200S, Leica Biosystems, Buffalo Grove, IL). Brain slices (230 µm) containing either the KF, BötC, preBötC, or rVRG (identified based on anatomical landmarks and coordinates from Franklin and Paxinos, 2008) were prepared in artificial cerebrospinal fluid (aCSF) that contained the following (in mM): 126 NaCl, 2.5 KCl, 1.2 MgCl2, 2.4 CaCl2, 1.2 NaH2PO4, 11 d-glucose and 21.4 NaHCO3 (equilibrated with 95% O2– 5% CO2). Slices were stored at 32°C in glass vials with equilibrated aCSF. MK801 (10 µM) was added to the cutting and initial incubation solution (at least 30 minutes) to block NMDA receptor-mediated excitotoxicity. Brain slices were transferred to a recording chamber and perfused with 34°C aCSF (Warner Instruments, Hamden, CT) at a rate of 1.5–3 ml min^−1^.

Cells were visualized using an upright microscope (Nikon FN1) equipped with custom built LED-based IR-Dodt gradient contrast illumination and DAGE-MTI IR1000 camera. Cells containing FluoSpheres (580/605) or tdTomato were identified using LED epifluorescence illumination and a Texas Red filter cube (ex 559 nm/ em 630 nm). Whole-cell recordings were made using a Multiclamp 700B amplifier (Molecular Devices, Sunnyvale, CA) in voltage-clamp mode (Vhold = −60 mV). Glass recording pipettes (1.5-3 MΩ) were filled with internal solution that contained (in mM): 115 potassium methanesulfonate, 20 NaCl, 1.5 MgCl2, 5 HEPES(K), 2 BAPTA, 1–2 Mg-ATP, 0.2 Na-GTP, adjusted to pH 7.35 and 275–285 mOsM. The liquid junction potential (10 mV) was not corrected. Data were low-pass filtered at 10 kHz and collected at 20 kHz with pCLAMP 10.7 (Molecular Devices, Sunnyvale, CA), or collected at 400 Hz with PowerLab (LabChart version 5.4.2; AD Instruments, Colorado Springs, CO). Series resistance was monitored without compensation and remained <15 MΩ for inclusion. For optogenetic experiments, ChR2-expressing KF terminals were stimulated using 470 nm LED illumination (5 ms duration) through a 40x objective to optogenetically-evoke excitatory postsynaptic currents (oEPSC) in preBötC and rVRG neurons. A pair of optical stimuli (5 ms pulse, 50 ms interval) was delivered every 20 seconds. Blockers of glycine (strychnine, 1 µM) and GABA-A (picrotoxin, 100 µM) receptors were added to the aCSF to isolate excitatory neurotransmission. Peak amplitudes were determined in Clampfit 10.7 (Molecular Devices), and paired-pulse ratios (peak 2/peak 1), were determined in Microsoft Excel. All drugs were applied by bath perfusion. Bestatin (10 µM) and thiorphan (1 µM) were included with ME to prevent degradation.

### Immunohistochemistry and microscopy

Mice were anesthetized with isoflurane and transcardially perfused with phosphate-buffered saline (PBS) followed by 10% formalin. The brains were removed and stored at 4°C in cryoprotectant (30% sucrose in 10% formalin). A vibratome (VT 1200S, Leica Biosystems, Buffalo Grove, IL) was used to prepare free-floating coronal brain slices (40-100 µm) for microscopy or immunohistochemistry.

Free-floating slices were stained for forkhead box P2 (FoxP2), calcitonin gene-related peptide (CGRP), or neurokinin 1 receptor (NK1R). Slices were washed in diluting buffer (TBS with 2% bovine serum albumin, 0.4% Triton X-100, and 1% filtered normal goat serum) for 30 min, blocked in TBS and 20% normal donkey serum for 30 minutes and incubated in primary antibody for 24 hours at 4°C. Primary antibodies included: sheep polyclonal anti-FoxP2 (cat.# AF5647; R&D Systems, Minneapolis, MN; 1:1000 in diluting buffer), rabbit polyclonal anti-CGRP (cat.# T-4032; Peninsula, San Carlos, CA, 1:1000 in diluting buffer, rabbit polyclonal anti-NK1R (cat.#S8305; Sigma-Aldrich; 1:1000 in diluting buffer). Slices were washed in diluting buffer and then incubated in secondary antibody (goat anti-rabbit 647 (cat. # A32733; Thermo Fisher Scientific, Waltham, MA) or donkey anti-sheep 647 (cat. # A21448; Thermo Fisher Scientific; 1:500)) in diluting buffer. Finally, slices were rinsed with TBS and ddH_2_0 and mounted onto glass slides with Fluoromount-G DAPI (Thermo Fisher Scientific). A confocal laser scanning microscope (Nikon A1R) with a ×10 objective (N.A. 0.3) or a multizoom microscope (Nikon AZ100) with a ×1 objective (N.A. 0.1) were used to image sections. All images were processed in Fiji (Schindelin *et al*., 2012).

To determine the spread and intensity of mCherry expression in the BötC, preBötC, and rVRG serial coronal brain slices (50 µm) were collected and every slice containing mCherry expression was imaged in sequential order with a multizoom microscope (Nikon AZ100) at 500 ms exposure. Mean fluorescence intensity was determined for a region of interest drawn ventral to the NA to encompass the BötC, preBötC, or rVRG in sequential slices. Mean intensity data were background subtracted and normalized to the peak intensity per injection. Bregma level was assigned using anatomical landmarks, including the inferior olives, nucleus ambiguus, and nucleus tractus solitarius (Franklin & Paxinos, 2008; Varga *et al*., 2020).

### Drugs

ME ([Met^5^]-enkephalin acetate salt), bestatin, DL-thiorphan, strychnine, picrotoxin, DNQX, 4-aminopyridine (4AP) and MK801 were from Sigma-Aldrich (St Louis, MO, USA). Tetrodotoxin and ML-297 was from Tocris Bio-Techne (Minneapolis, MN).

### Statistics

All statistical analyses were performed in GraphPad Prism 8 (La Jolla, CA). All error bars represent SEM unless otherwise stated. Replicates are biological replicates. Data with n > 8 were tested for normality with Kolmogorov-Smirnov tests. Comparisons between two groups were made using paired or unpaired two-tailed t-tests. Comparisons between three or more groups were made using one-way ANOVA followed by Tukey’s post-hoc test.

**Table 1.**
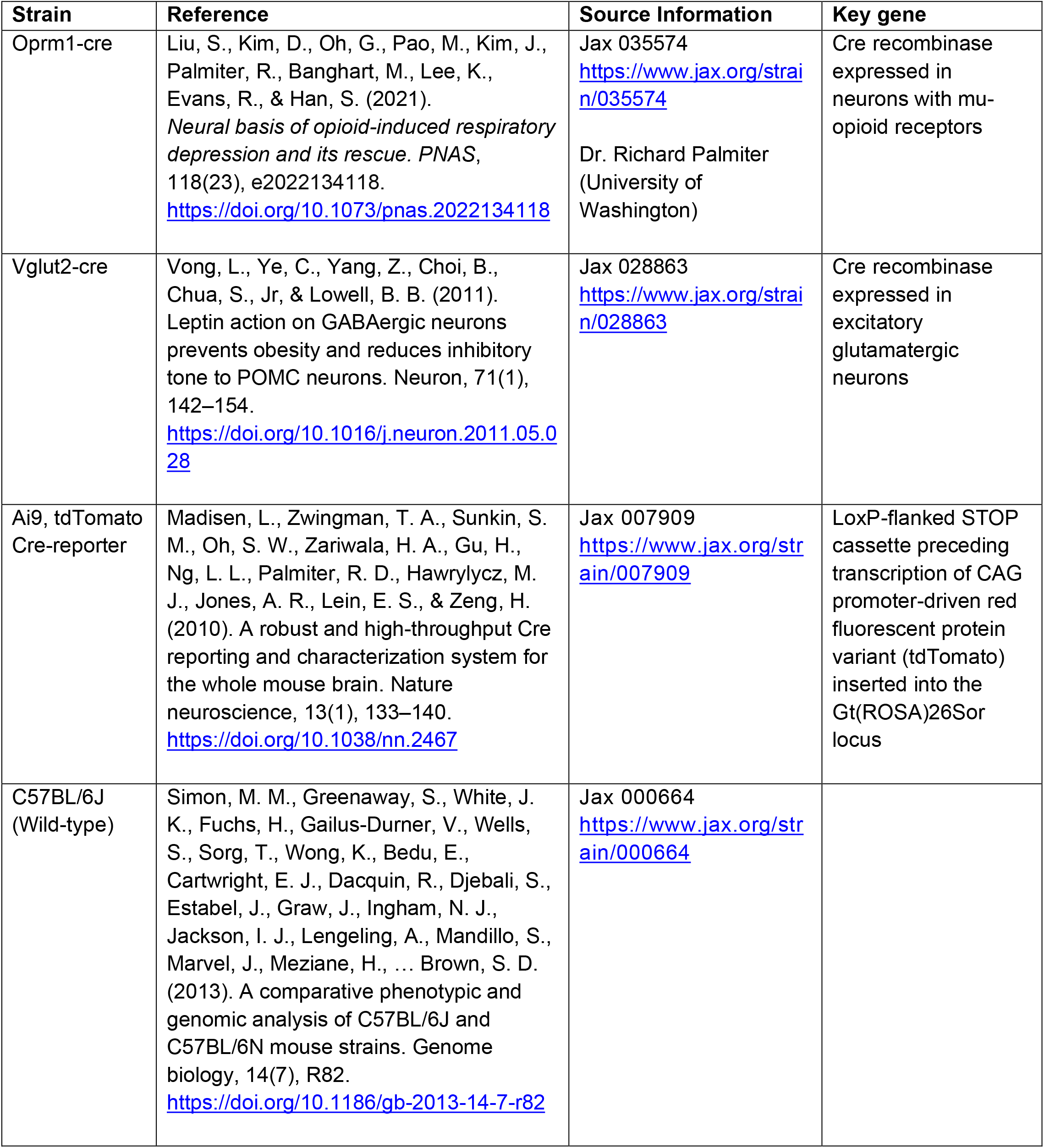

| Strain | Reference | Source Information | Key gene |
| --- | --- | --- | --- |
| Oprm1-cre | Liu, S., Kim, D., Oh, G., Pao, M., Kim, J., Palmiter, R., Banghart, M., Lee, K., Evans, R., & Han, S. (2021). *Neural basis of opioid-induced respiratory depression and its rescue. PNAS*, 118(23), e2022134118. https://doi.org/10.1073/pnas.2022134118 | Jax 035574<br>https://www.jax.org/strain/035574<br>Dr. Richard Palmiter<br>(University of Washington) | Cre recombinase expressed in neurons with mu-opioid receptors |
| Vglut2-cre | Vong, L., Ye, C., Yang, Z., Choi, B., Chua, S., Jr., & Lowell, B. B. (2011). Leptin action on GABAergic neurons prevents obesity and reduces inhibitory tone to POMC neurons. *Neuron*, 71(1), 142–154. https://doi.org/10.1016/j.neuron.2011.05.028 | Jax 028863<br>https://www.jax.org/strain/028863 | Cre recombinase expressed in excitatory glutamatergic neurons |
| Ai9, tdTomato Cre-reporter | Madisen, L., Zwingman, T. A., Sunkin, S. M., Oh, S. W., Zariwala, H. A., Gu, H., Ng, L. L., Palmiter, R. D., Hawrylycz, M. J., Jones, A. R., Lein, E. S., & Zeng, H. (2010). A robust and high-throughput Cre reporting and characterization system for the whole mouse brain. *Nature neuroscience*, 13(1), 133–140. https://doi.org/10.1038/nn.2467 | Jax 007909<br>https://www.jax.org/strain/007909 | LoxP-flanked STOP cassette preceding transcription of CAG promoter-driven red fluorescent protein variant (tdTomato) inserted into the Gt(ROSA)26Sor locus |
| C57BL/6J (Wild-type) | Simon, M. M., Greenaway, S., White, J. K., Fuchs, H., Gailus-Durner, V., Wells, S., Sorg, T., Wong, K., Bedu, E., Cartwright, E. J., Dacquin, R., Djebali, S., Estabel, J., Graw, J., Ingham, N. J., Jackson, I. J., Lengeling, A., Mandillo, S., Marvel, J., Meziane, H., ... Brown, S. D. (2013). A comparative phenotypic and genomic analysis of C57BL/6J and C57BL/6N mouse strains. *Genome biology*, 14(7), R82. https://doi.org/10.1186/gb-2013-14-7-r82 | Jax 000664<br>https://www.jax.org/strain/000664 |  |

**Table 2.**
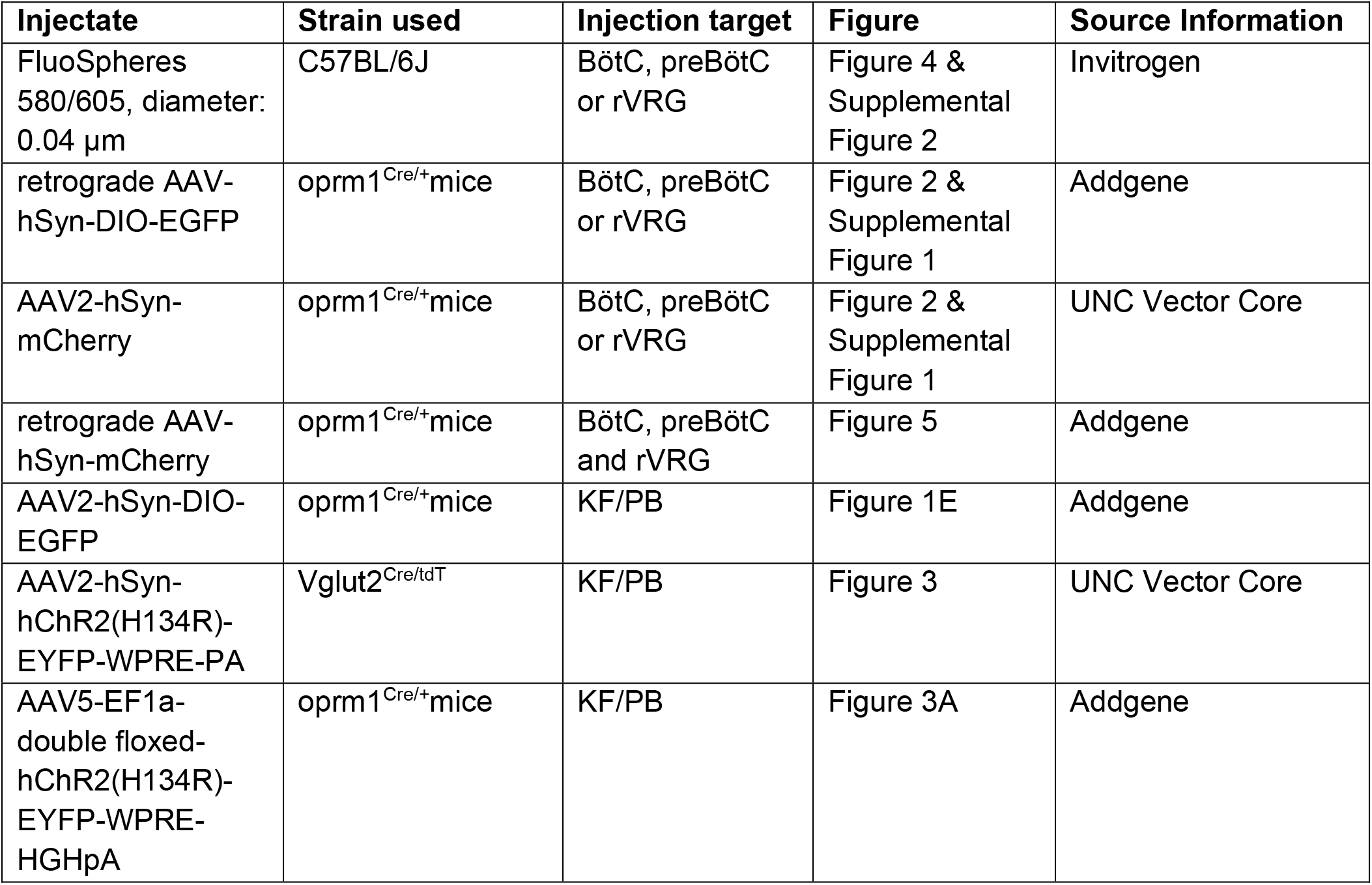

| **Injectate** | **Strain used** | **Injection target** | **Figure** | **Source Information** |
| --- | --- | --- | --- | --- |
| FluoSpheres 580/605, diameter: 0.04 μm | C57BL/6J | BötC, preBötC or rVRG | Figure 4 & Supplemental Figure 2 | Invitrogen |
| retrograde AAV-hSyn-DIO-EGFP | oprm1<sup>Cre/+</sup> mice | BötC, preBötC or rVRG | Figure 2 & Supplemental Figure 1 | Addgene |
| AAV2-hSyn-mCherry | oprm1<sup>Cre/+</sup> mice | BötC, preBötC or rVRG | Figure 2 & Supplemental Figure 1 | UNC Vector Core |
| retrograde AAV-hSyn-mCherry | oprm1<sup>Cre/+</sup> mice | BötC, preBötC and rVRG | Figure 5 | Addgene |
| AAV2-hSyn-DIO-EGFP | oprm1<sup>Cre/+</sup> mice | KF/PB | Figure 1E | Addgene |
| AAV2-hSyn-hChR2(H134R)-EYFP-WPRE-PA | Vglut2<sup>Cre/tdT</sup> | KF/PB | Figure 3 | UNC Vector Core |
| AAV5-EF1a-double floxed-hChR2(H134R)-EYFP-WPRE-HGHpA | oprm1<sup>Cre/+</sup> mice | KF/PB | Figure 3A | Addgene |

**Table 3.**

| **Antigen** | **Immunogen description** | **Source, host species, RRID** | **Concentration** |
| --- | --- | --- | --- |
| Forkhead box P2 (FoxP2) | Targets human and mouse FoxP2 | R&D Systems, sheep polyclonal, cat.# AF5647, RRID: AB_2107133 | 1:1000 |
| Calcitonin gene-related peptide (CGRP) | Targets alpha-CGRP in canine, mouse, and rat | Peninsula, rabbit polyclonal, cat.# T-4032, RRID: AB_518147 | 1:1000 |
| Neurokinin 1 receptor (NK1R) | Targets C-terminal of NK1R in mouse, guinea pig, and human | Sigma-Aldrich, rabbit polyclonal, cat.# S8305 RRID: AB_261562 | 1:1000 |

## Acknowledgements

This work was supported by National Institutes of Health Grant R01DA047978 (ESL). JTB was supported by F31DA053798. We would like to thank Dr. Richard Palmiter (University of Washington) for generously providing the Oprm1-Cre mice. We thank Drs. Gordon Mitchell, John Williams and Adrienn Varga for comments on the manuscript.

## Author contributions

ESL designed research. JTB performed experiments and analyzed data. JTB prepared figures and drafted the manuscript. JTB and ESL revised and approved the final version of the manuscript.

## Additional Information

The authors declare no competing interests, financial or otherwise.

## Data Availability

Data generated or analyzed during this study are included in the manuscript and supporting files.

**Figure 2 – Figure supplement 1.**
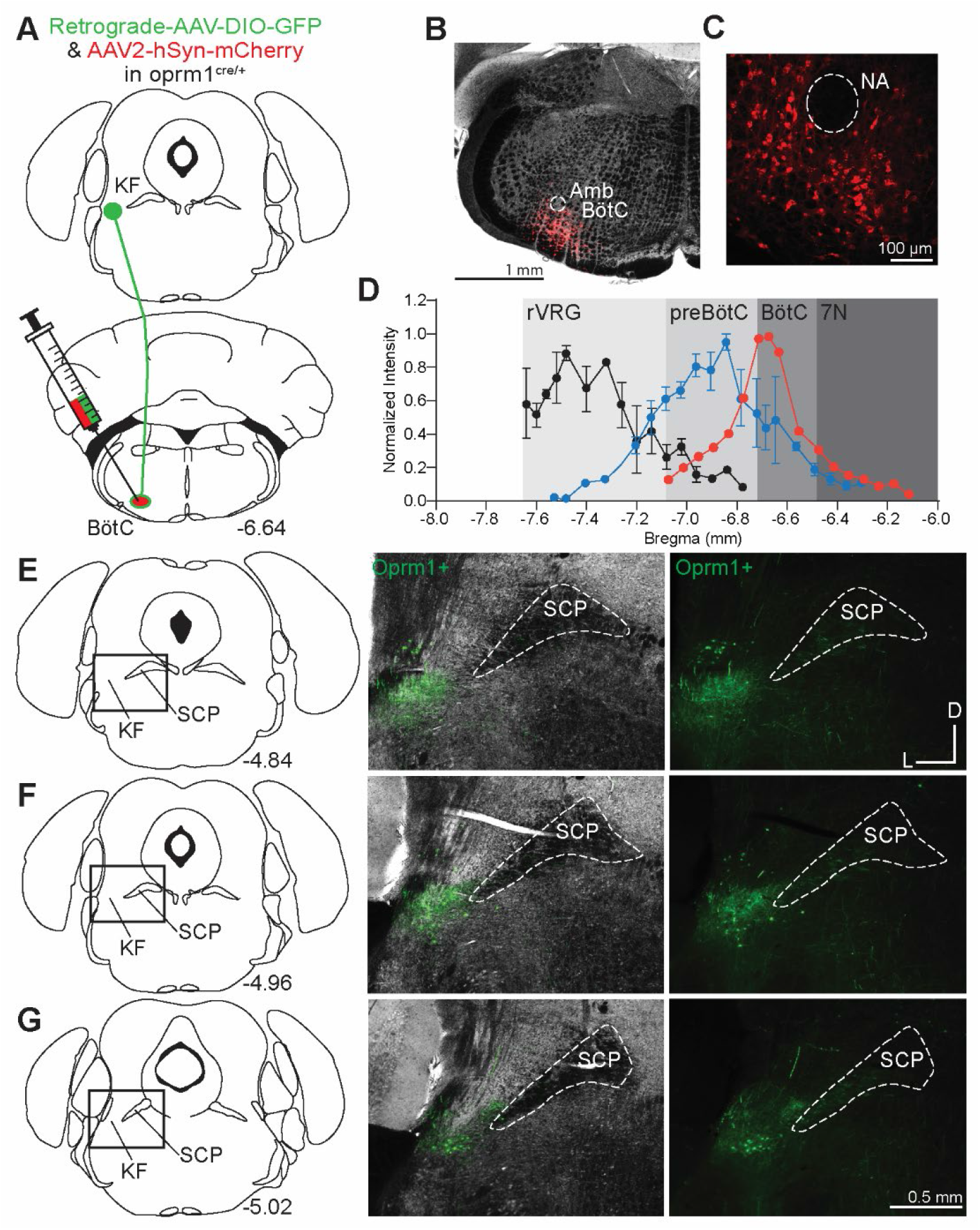
Oprm1+ KF neurons project to the BötC. **(A**) Schematic illustrating the approach to retrogradely label Oprm1+ KF neurons projecting to the BötC. **(B-C)** Images of coronal slices from the medulla with a control injection of AAV2-hSyn-mCherry into the BötC of an Oprm1^Cre/+^ mouse to mark injection spread. The nucleus ambiguous (NA) was used as a medullary marker in both images. **(D)** Quantification of normalized AAV2-hSyn-mCherry spread along the rostral to caudal axis in the ventrolateral medulla of representative injection into BötC. Data from injections into preBötC (n=5), and rVRG (n=5) are duplicated from Figure 2 for comparison. **(E-G)** Images of Oprm1+ expression following a BötC injection across three levels of the KF/LPB. The bregma coordinates and approximate location of KF/LPB are indicated. The scale bar in (G) applies to all images in E-G.

**Figure 4 – Figure supplement 1.**
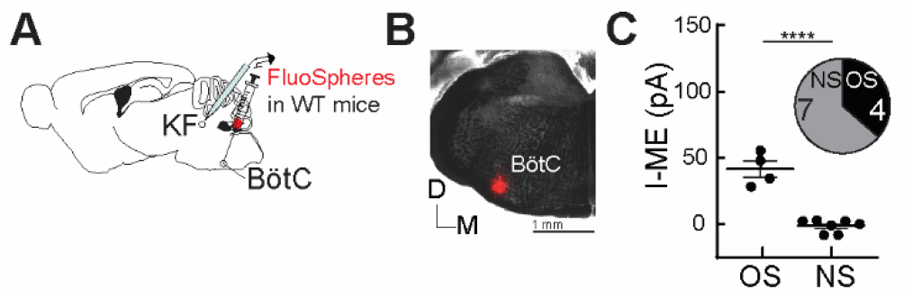
Opioids hyperpolarize KF neurons that project to the BötC. **(A)** Schematic of approach to retrogradely label KF neurons that project to the BötC with FluoSpheres in wild-type mice. **(B)** Image of FluoSpheres injection into the BötC. **(C)** Quantification of ME-mediated current in opioid-sensitive (OS) and non-opioid-sensitive (NS) KF neurons that project to the BötC (n=11; p=0.0001; unpaired t-test). ME (1 µM) induced an outward current in 4 of 11 KF neurons that project to the BötC. Individual data points are from individual neurons in separate slices. Line and error are mean ± SEM.

**Figure 5 – Figure supplement 1.**
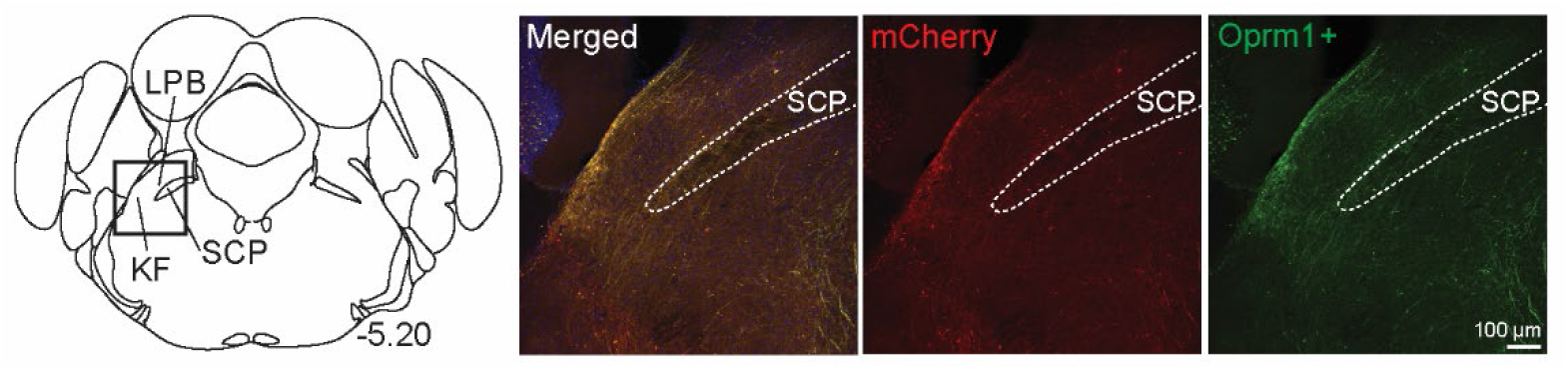
Medullary projecting Oprm1+ neurons are mostly absent from the caudal KF and lateral parabrachial areas. **(A)** Retrograde labeled neurons (both Oprm1+ and Oprm1-) were mostly lacking in caudal KF or lateral parabrachial area. The bregma coordinate and approximate location in KF/LPB are indicated. The scale bar applies to all images. KF (Kölliker-Fuse) and (LPB) lateral parabrachial.

